# Oxidation-sensitive cysteines drive IL-38 amyloid formation

**DOI:** 10.1101/2023.03.26.534252

**Authors:** Alejandro Diaz-Barreiro, Gea Cereghetti, Jenna Tonacini, Dominique Talabot-Ayer, Sylvie Kieffer-Jaquinod, Arnaud Huard, Christopher Swale, Yohann Couté, Matthias Peter, Antonio Francés-Monerris, Gaby Palmer

**Affiliations:** Division of Rheumatology, Department of Medicine, Faculty of Medicine, University of Geneva, Geneva, Switzerland; Department of Pathology and Immunology, Faculty of Medicine, University of Geneva, Geneva, Switzerland; Geneva Centre for Inflammation Research, Geneva, Switzerland; Department of Chemistry, University of Cambridge, Cambridge, United Kingdom; Université Grenoble Alpes, CEA, INSERM, UA13 BGE, CNRS, CEA, FR2048, Grenoble, France; Institute for Advanced Biosciences (IAB), Team Host-Pathogen Interactions and Immunity to Infection, University Grenoble Alpes, INSERM U1209, CNRS UMR5309, Grenoble, France; Institute of Biochemistry, Department of Biology, ETH Zurich, Zurich, Switzerland; Institut de Ciència Molecular, Universitat de València, 46071 València, Spain

## Abstract

Cytokines of the interleukin (IL)-1 family are widely expressed in epithelial surfaces, including the epidermis, where they play a key role in the maintenance of barrier integrity and host defense. A recent report associated the IL-1 family member IL-33 with stress granules (SGs) in epithelial cells. Formation of SGs is promoted by the aggregation of proteins harboring low complexity regions (LCRs). In this study, using computational analyses, we predicted the presence of LCRs in six of the eleven IL-1 family members. Among these, IL-38 contained a long LCR and localized to Ras GTPase-activating protein binding protein 1 (G3BP1) positive SGs, as well as to G3BP1 negative intracellular protein condensates in keratinocytes exposed to oxidative stress (OS). In addition, we identified two highly aggregation-prone amyloid core (AC) motifs in the IL-38 LCR and detected the formation of amyloid IL-38 aggregates in response to OS in cells and *in vitro*. Disulfide bond mapping, *in silico* modelling and the analysis of specific cysteine mutants supported a model in which specific oxidation-sensitive cysteines act as redox switches to modify the conformation of IL-38 and thus the surface exposure of its ACs, shuttling it from a soluble state into biomolecular condensates. Finally, the presence of IL-38 granules in human epidermal layers highly exposed to environmental OS suggests that oxidation-induced formation of amyloid aggregates, as a previously unrecognized intrinsic biological property of IL-38, may be physiologically relevant at this epithelial barrier.

## Introduction

The interleukin (IL)-1 family of cytokines includes 7 pro-inflammatory agonists, namely IL-1α, IL-1β, IL-18, IL-33, IL-36α, IL-36β and IL-36γ, and 4 anti-inflammatory members, IL-1Ra, IL-36Ra, IL-37 and IL-38. These cytokines are widely expressed in epithelia, where they play an essential role in the maintenance of barrier integrity and host defense (Matarazzo et al., 2022). Interestingly, the 4 anti-inflammatory family members are constitutively produced and accumulate at steady state in epidermal keratinocytes (Martin et al., 2021). This may reflect their role in preserving skin immune homeostasis and tolerance, but also led to the suggestion that they might exert additional functions in the epidermis, unrelated to inflammation (Lachner et al., 2017).

Despite the importance of their extracellular activity in the control of inflammatory responses, most IL-1 family cytokines lack a signal peptide for classical ER-Golgi secretion and rely on unconventional secretion mechanisms or passive release upon cell membrane disruption to relocate to the extracellular space. In addition, for some family members, intracellular effects have been described (Banda et al., 2005; Carriere et al., 2007; Nold et al., 2010; Palmer et al., 2005; Roussel et al., 2008). In particular, IL-1α and IL-33 are considered as dual function molecules, which exert a physiological role inside their producing cells, but also act as extracellular alarmins when they are released from damaged, stressed or activated cells (Bertheloot and Latz, 2017). In this context, the release of IL-33 from epithelial cells exposed to allergen proteases was recently reported to involve its association with stress granules (SGs) (Chen et al., 2022).

SGs are membraneless organelles that are formed by liquid-liquid phase separation. Formation of these biomolecular condensates is promoted by the aggregation of proteins harboring low complexity regions (LCRs) (Cereghetti et al., 2018; Guillen-Boixet et al., 2020), which led us to examine the presence of LCRs in IL-1 family cytokines. We identified a LCR including two amyloid core (AC) sequences in the human IL-38 protein and detected the formation of amyloid IL-38 aggregates in oxidative conditions, in cells and *in vitro.* Disulfide bond mapping, *in silico* simulations, and the analysis of specific cysteine mutants led us to propose a molecular model in which disulfide bond formation between two vicinal cysteines leads to increased AC surface exposition, thereby promoting IL-38 aggregation. Remarkably, we observed similar granular IL-38 structures in cultured keratinocytes during OS and in healthy human epidermis *ex vivo,* suggesting that the formation of IL-38 condensates may be involved in epidermal differentiation and skin homeostasis.

## Results

### IL-38 forms condensates in keratinocytes exposed to oxidative stress

Recent findings revealed that upon cellular stress IL-33 localizes to stress granules (SGs) (Chen *et al.,* 2022). To investigate whether other IL-1 family members might display similar tendencies to condense and be recruited to SGs, we analyzed their protein sequence using computational tools to identify putative aggregation-prone regions (**Fig 1a**) (Wootton, 1994). We found that IL-33 and IL-38 harbor the longest low complexity regions (LCRs), encompassing amino acids (aa) 138-166 and 95-122, respectively. In contrast, other IL-1 family members contain no aggregation-prone sequence (IL-36β, IL-36γ, IL-18, IL-1Ra, and IL-36Ra) or shorter stretches, less likely to trigger aggregation (IL-1α, IL-1β, IL-36α and IL-37) (**Fig 1a**). Since LCRs are thought to play fundamental roles in driving the formation of protein condensates (Martin and Mittag, 2018), we hypothesized that IL-38, like IL-33, might form protein condensates in stressed cells. IL-38 is constitutively expressed by keratinocytes in the outermost layer of the skin, which is continuously exposed to environmental threats causing oxidative stress (OS). We thus used a pelleting assay to examine IL-38 aggregation in a human keratinocyte cell line (NHK/38) exposed to oxidation. To this end, NHK/38 cells were treated (Men) or not (NT) with the oxidant menadione and IL-38 solubility followed by immunoblotting (**Fig 1b**). Strikingly, while IL-38 was mostly soluble in the cytoplasm of untreated control cells, it was markedly re-localized to an insoluble pellet fraction in extracts prepared from menadione-treated cells.

**Figure 1.**
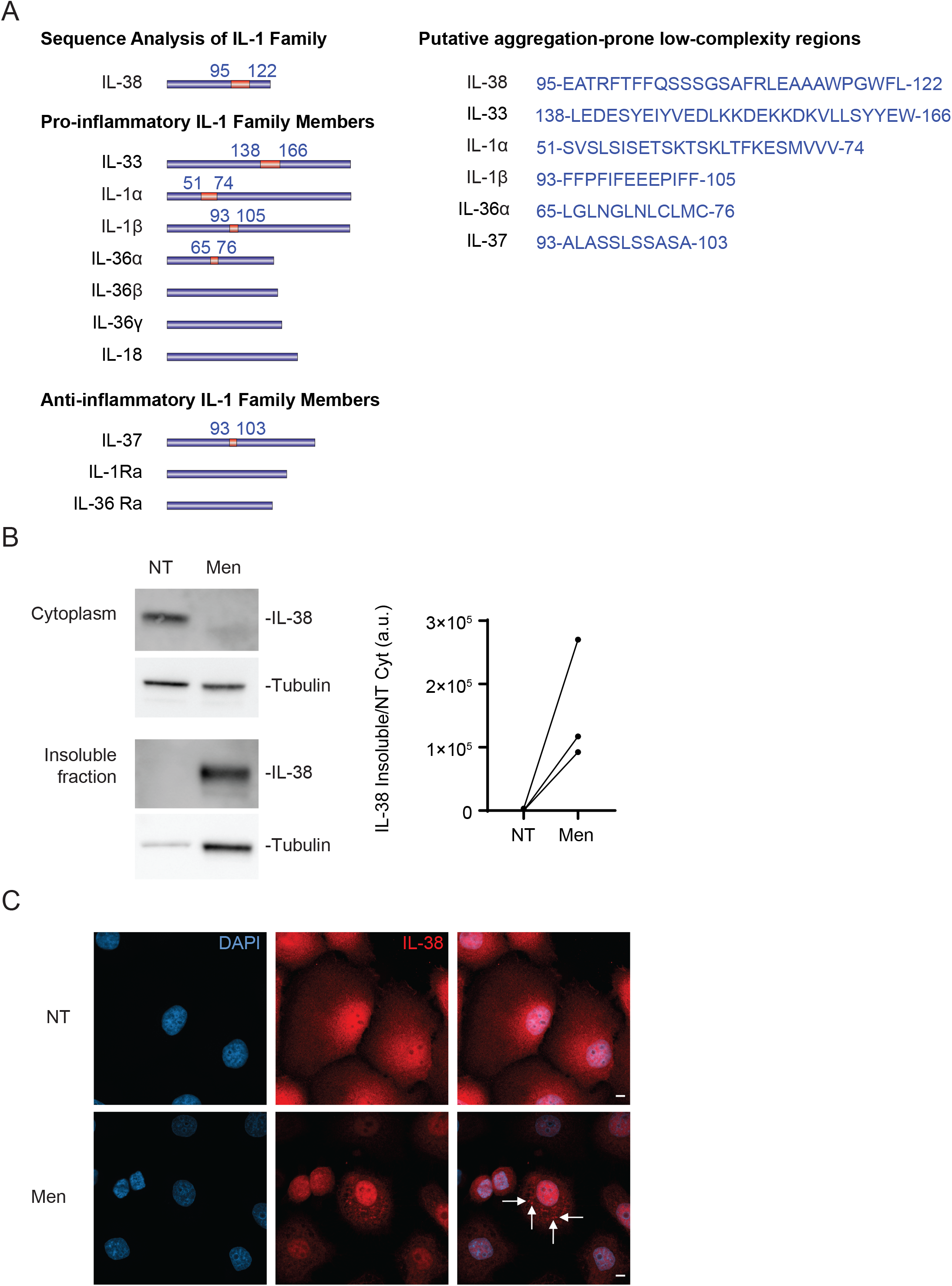
IL-38 contains a LCR and forms condensates in keratinocytes during OS. (A) Schematic representation (orange, left panel) and sequences (right panel) of LCRs identified in the major IL-1 family cytokine isoforms. IL-33 and IL-38 harbor the longest predicted LCRs. (B) IL-38 levels were assessed in the soluble cytoplasmic fraction (left, top panels), and in the insoluble fraction (left, bottom panels) by reducing WB in Dox-induced NHK/38 cells treated (Men), or not (NT), with 75 μM menadione for 1 h. Membranes were stripped and reblotted for tubulin. ImageJ densitometric quantification (right panel) of WB bands represents the amount of IL-38 in the insoluble fraction with (Men) or without (NT) menadione-treatment, normalized against IL-38 levels in the cytoplasmic fraction in NT cells. Results are shown as paired values, each line representing one out of n=3 independent experiments. a.u., arbitrary units. (C) Localization of IL-38 (red staining; second and third columns) was examined by IF and confocal microscopy in Dox-induced NHK/38 cells treated with 75 μM menadione (Men) or its vehicle (NT) for 1 h. IL-38 formed granular aggregates upon Men treatment (white arrows). DAPI (blue staining; first and third columns) was used to stain the nuclei. Scale bars: 5 μm.

To corroborate these results, immunofluorescence microscopy (IF) was performed to visualize IL-38 aggregation in NHK/38 cells. Interestingly, IL-38-positive granules were observed in menadione-treated NHK/38 cells (**Fig 1c, lower panels, white arrows**), while the staining was more homogenously distributed in untreated controls (**Fig 1c, top panels**). IL-38 in the soluble cytoplasmic fraction decreased upon menadione treatment (**Fig 1b, top panels**). This decrease was not explained by disrupted plasma membrane integrity, as illustrated by the unaltered levels of the cytoplasmic enzyme LDH in the culture media after 2 hours of OS (**Supp Fig 1a**). Analysis of p62 (**Supp Fig 1b**) or cleaved caspase 3 (**Supp Fig 1c**) excluded the formation of autophagosomes or apoptotic bodies in menadione-treated NHK/38 cells. However, staining of the Ras GTPase-activating protein binding protein 1 (G3BP1) indicated that some IL-38 aggregates co-localize with SGs (**Supp Fig 1d, top panels, solid arrows**), while others predominantly or exclusively stained for IL-38 (**Supp Fig 1d, top panels, dashed arrows**). To better visualize OS-regulated IL-38 foci, soluble material was extracted from permeabilized cells before IF staining (**Supp Fig 1d, bottom panels**). While cytoplasmic extraction caused loss of IL-38 staining in control cells, G3BP1-positive and negative IL-38 foci remained detectable in menadione-treated NHK/38 cells (**Supp Fig 1d, bottom panels, solid and dashed arrows, respectively**).

Taken together, these data indicate that IL-38 contains a predicted aggregation-prone sequence (aa 93-122), and localizes to G3BP1 positive and negative intracellular protein condensates in keratinocytes exposed to OS.

### IL-38 forms amyloid-like structures in oxidative conditions in vitro and in cells

Further computational analysis of the IL-38 protein sequence revealed that its LCR contains two regions (aa 98-103 and 119-123) with high propensity to form amyloids, which we termed amyloid cores (ACs) (**Fig 2a**). Amyloids are a particular type of protein aggregates, characterized by a cross-beta secondary structure and the ability to resist SDS denaturation. We thus used semi-denaturing detergent agarose gel electrophoresis (SDD-AGE) (Bagriantsev et al., 2006) to compare the presence of amyloid structures in untreated or stressed HEK cells expressing IL-38 (**Fig 2b**). As expected, IL-38 migrated as a monomer in untreated controls, while a smear characteristic of amyloids appeared in H_2_O_2_-treated cells, strongly suggesting that IL-38 forms amyloids in response to OS.

**Figure 2.**
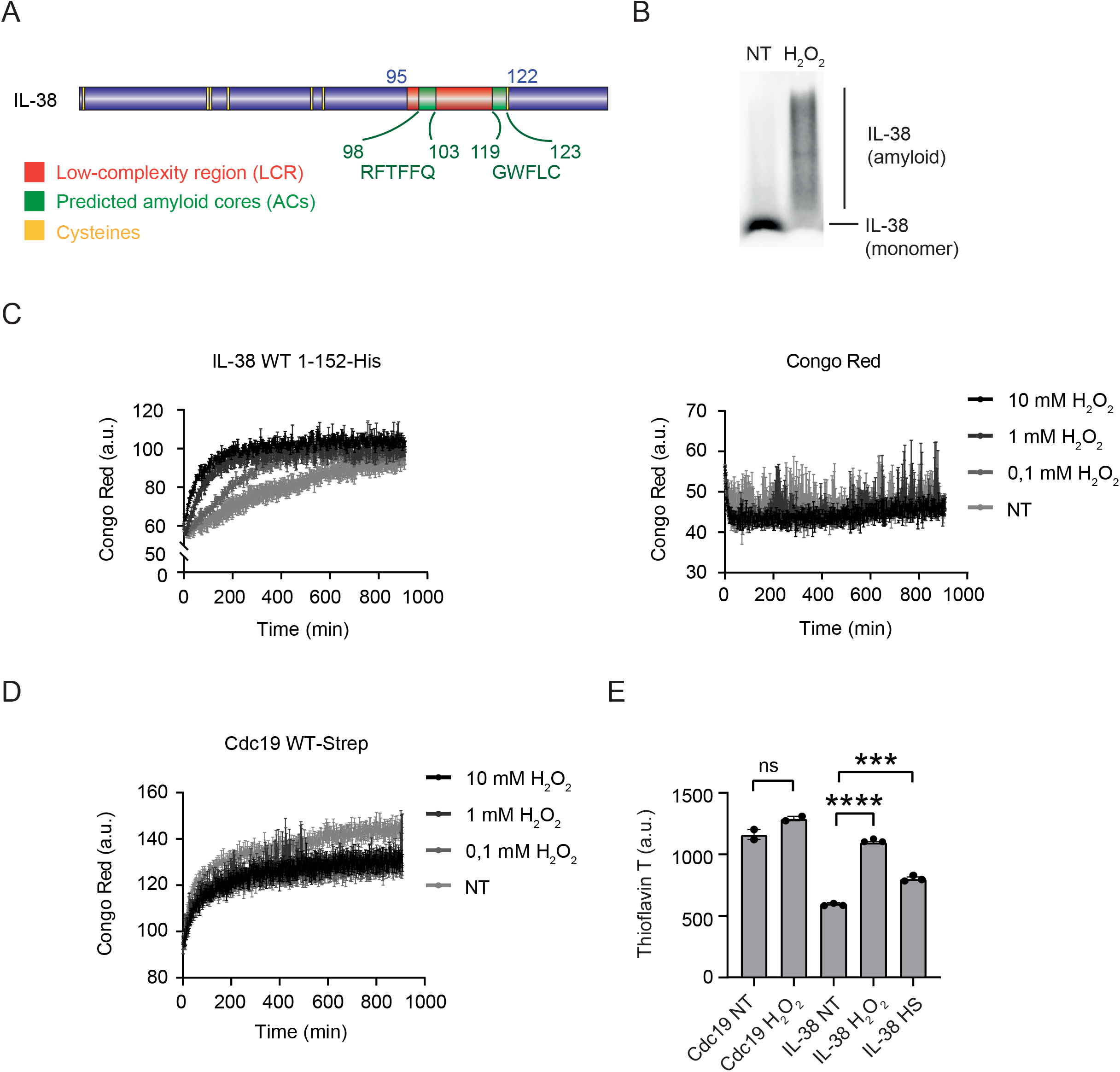
IL-38 forms amyloid structures *in vitro* and in cells in oxidizing conditions. (A) Schematic representation of the IL-38 sequence highlighting the predicted LCR (orange), putative ACs (green, aa 98-103 and 119-123), and cysteines (yellow). (B) The formation of intracellular IL-38 amyloid structures was examined by SDD-AGE and WB in HEK cells transfected with pcDNA4/TO/hIL-38-TD and treated or not (NT) with H_2_O_2_ (10 mM) for 1 h. The positions of IL-38 monomer (NT, band) and amyloids (H_2_O_2_, smear) are indicated. Results are representative of n=3 independent experiments. (C) Purified C-terminally His-tagged aa 1-152 recombinant human IL-38 protein (10 μM, left panel) was exposed to H_2_O_2_ (0.1, 1 and 10 mM, grey to black symbols), or left untreated (NT, lightest grey), and the formation of IL-38 amyloid structures was assessed by CR staining. CR alone (right panel) was equally exposed to H_2_O_2_ (0.1, 1 and 10 mM, grey to black symbols), or left NT (lightest grey) at 37 °C, in order to exclude unspecific detection of CR fluorescence. (D) Streptavidin-tagged recombinant Cdc19 protein, used as a specificity control, was exposed to H_2_O_2_ (0.1, 1 and 10 mM, grey to black symbols), or left NT (lightest grey), and the *in vitro* formation of Cdc19 amyloids was assessed by CR staining. (c-d) Results are shown as individual values and means + SEM for n=3 independent experiments. (E) C-terminally His-tagged full-length recombinant IL-38 and streptavidin-tagged Cdc19 were exposed to 100 mM H_2_O_2_ or left NT for 30 min on ice. IL-38 was also subjected to a heat shock (HS) at 45 °C for 30 min and ThT fluorescence was measured. Results are shown as individual values and means + SEM for n=3 (IL-38) or n=2 (Cdc19) independent experiments. ***p<0,001, ****p<0,0001, by unpaired t test. A.u., arbitrary units. WT, wild type.

To characterize IL-38 aggregates *in vitro,* we tested the amyloidogenic properties of a C-terminally His-tagged, full-length human IL-38 protein in oxidative conditions. Specifically, we measured its staining with Congo Red (CR), a dye that emits red fluorescence upon binding to amyloid fibrils (**Fig 2c, left panel**). Interestingly, addition of H_2_O_2_ increased CR fluorescence in a dose-dependent manner. At 10 mM H_2_O_2_, CR staining increased rapidly after H_2_O_2_ addition, reaching already 50% of the maximal effect after 40 min. H_2_O_2_ *per se* had no effect on CR fluorescence at any of the tested concentrations (**Fig 2c, right panel)**. Likewise, H_2_O_2_ (0.1-10 mM) failed to enhance but rather slightly decreased CR-binding, and thus amyloid assembly, of the pH-regulated, reversible yeast amyloid Cdc19 (**Fig 2d**), highlighting the specificity of IL-38 amyloid formation regulated by OS. Staining of IL-38 with another amyloid-specific dye, Thioflavin T (ThT) yielded similar results (**Fig 2c, Supp Fig 2**). Next, we assessed the ability of heat shock, as another stressor, to induce IL-38 amyloid formation *in vitro* by measuring ThT staining. While heat shock increased IL-38 ThT staining, this effect was less pronounced compared to incubation with 100 mM H_2_O_2_ for 30 min (**Fig 2e**), suggesting that OS specifically and efficiently triggers the formation of IL-38 amyloids *in vitro.*

Altogether, these findings indicate that IL-38 contains two predicted AC motifs and forms amyloid structures in oxidative conditions both *in vitro* and in cells.

### Oxidation-sensitive cysteines act as redox switches to control IL-38 aggregation in vitro and in cells

Next, we further investigated the molecular mechanisms underlying oxidation-dependent IL-38 amyloid formation. In the IL-38 crystal (PDB: 5BOW) and in the AlphaFold-modelled IL-38 protein, the two ACs mostly fall into β-sheets buried inside the protein (**Fig 3a).** Intriguingly, although IL-38 has a remarkably high cysteine content (7 cysteines, amounting to 4.6% of its 152 aa sequence) (**Fig 2a**), the IL-38 crystal does not show any disulfide bonds. Based on the high structural homology within the IL-1 family and the presence of disulfide bridges in other family members (Cohen et al., 2015; Schreuder et al., 1997), we hypothesized that IL-38 might nevertheless be able to form intramolecular S-S bonds and measured the distances between the sulfurs of its 7 cysteine residues along a 1 μs accelerated molecular dynamics (aMD) *in silico* simulation (**Supp Fig 3a**). Based on the length of an equilibrium S-S bond (~2.05 to 3 Å), S-S distances from 3-3.5 to 4-4.5 Å, in which no solvent molecule can be placed between the two sulfur atoms, were considered as possible hotspots for S-S bond formation. This analysis identified two cysteine pairs, C2-C43 and C37-C38 as the only candidates. We then confirmed the presence of two S-S bonds in recombinant human IL-38 protein by top-down nanoLC-MS (**Supp Fig 3b**). In addition, nanoLC-MS/MS analysis of alkylated IL-38 revealed alkylation of C67, C70 and C123, whereas no MS/MS fragments were observed in the N-terminal part of the protein (**Supp Fig 3c**), strongly suggesting that the two disulfide bonds indeed involve C2, C37, C38 and C43, in agreement with our *in silico* predictions. Further *in silico* modelling predicted major and opposite effects of the C2-C43 and C37-C38 bonds on the global conformation of IL-38. The C2-C43 S-S bond is predicted to stabilize the region of the 12-β trefoil fold bearing the two ACs, and thus to keep them mostly hidden from the surface, even in presence of the C37-38 bond (**Supp Fig 3d**). In contrast, the C37-C38 S-S bond creates molecular tensions that disrupt the 12-β trefoil fold and strongly increase the AC surface exposure (**Fig 3b**). Accordingly, the predicted solvent-accessible surface area (SASA) of the two ACs was markedly increased in C37-C38 disulfide bonded (DSB), as compared to 12-β trefoil folded or C37-C38+C2-C43 DSB IL-38 (**Supp Fig 3e**).

**Figure 3.**
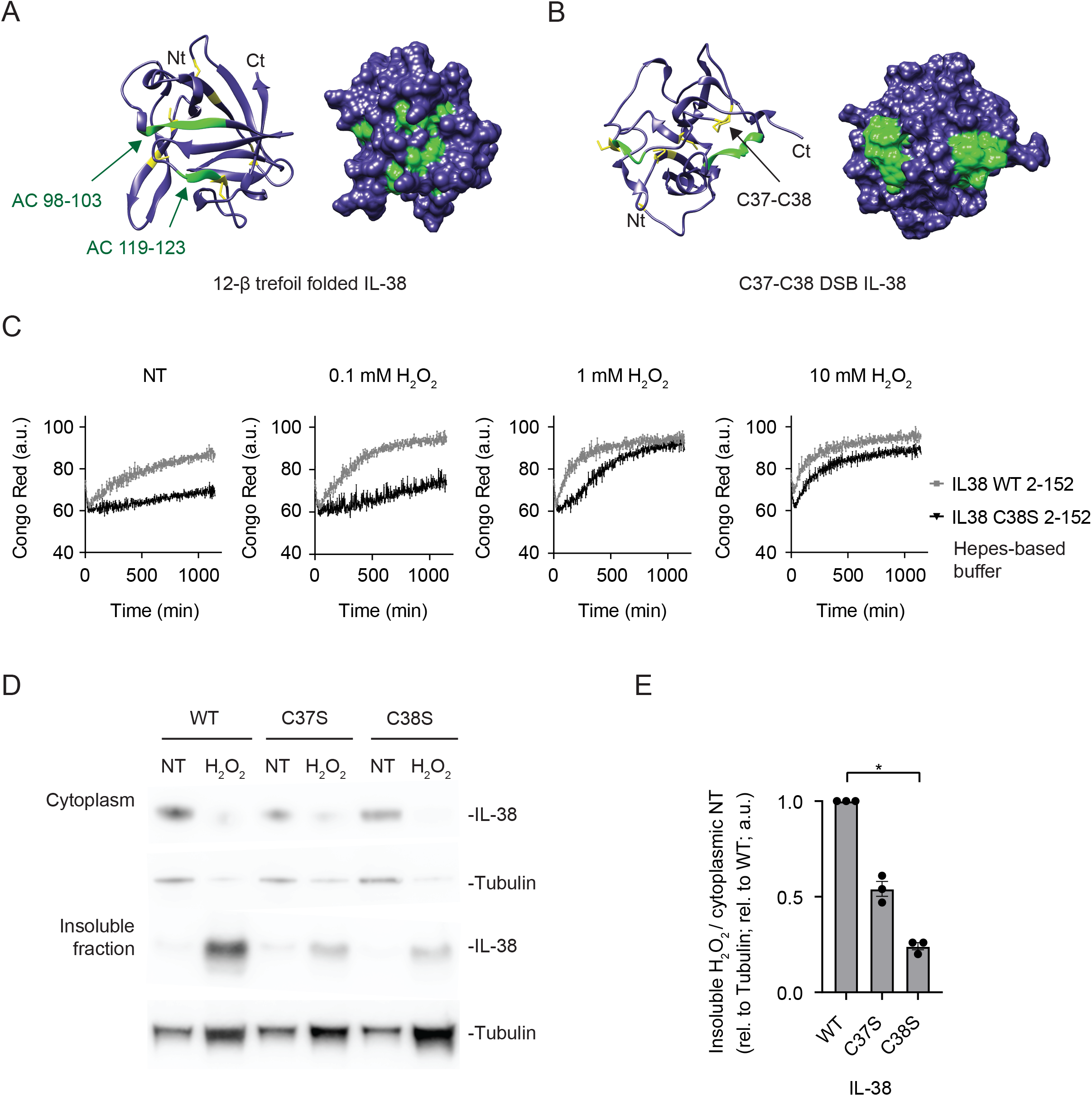
Oxidation sensitive cysteines act as redox switches to control IL-38 aggregation *in vitro* and in cells. (A) AlphaFold-modeled ribbon diagram (left) and surface representation (right) of the reduced IL-38 molecule without disulfide bridges, displaying the characteristic IL-1 family 12-β trefoil fold. (B) Ribbon diagram (left) and surface representation (right) of disulfide-bonded (DSB) IL-38 carrying a single C37-C38 disulfide bond (black arrow). The two ACs (green) are more exposed on the surface in (B). Cysteines are shown in yellow. (C) Recombinant WT (grey symbols) and C38S mutant (black symbols) human aa 2-152 IL-38 were treated with H_2_O_2_ (0.1, 1 and 10 mM), or left untreated (NT), and the *in vitro* formation of IL-38 amyloids was assessed by CR staining in a Hepes-based buffer. Results are expressed as mean + SEM for n=3 independent experiments. a.u., arbitrary units. (D) The levels of WT, C37S and C38S IL-38 were assessed by WB in the soluble cytoplasmic (top panels) and insoluble (bottom panels) fractions in transfected HEK cells treated or not (NT) with 10 mM H_2_O_2_ for 1 h. Membranes were stripped and reblotted for tubulin. (E) Densitometric quantification of WB bands associated to (D). Columns represent individual values and means + SEM of paired IL-38/tubulin ratios in the H_2_O_2_-treated insoluble fraction vs. the non-treated soluble cytoplasmic fractions for n=3 independent experiments. Results are shown as fold change relative to cells expressing WT IL-38. *p<0.05, by Friedman test with Dunn’s multiple comparisons. a.u., arbitrary units.

To experimentally link the C37-C38 bond to biochemical properties of IL-38, we generated a C38S mutant recombinant IL-38 protein, in which this bond cannot be formed, and compared the ability of WT and C38S IL-38 to form amyloid structures in the presence of increasing concentrations (0.1-10 mM) of H_2_O_2_ (**Fig 3c**). Strikingly, the C38S mutant showed a markedly decreased sensitivity to H_2_O_2_ oxidation, and significantly slower aggregation compared to WT IL-38 in presence of 0, 0.1 and 1 mM H_2_O_2_ (**Fig. 3c; Supp Fig 3f**). It is noteworthy that amyloid formation occurred with similar kinetics for WT and C38S IL-38 at the highest H_2_O_2_ concentration (10 mM, **Fig 3c; Supp Fig 3f**), indicating that the C38S mutation did not denature the protein or globally impair its aggregation properties. Altogether, *in silico* and *in vitro* findings suggest that oxidation of cysteine 38 enhances exposure of the aggregation-prone ACs, thus promoting IL-38 amyloid formation.

To test whether this mechanism is relevant also in the context of living cells, we then exposed WT, C37S or C38S IL-38 expressing HEK cells to OS. Similar to our observations in NHK/38 cells (**Fig 1c**), the three IL-38 proteins re-localized to the insoluble fraction in HEK cells upon H_2_O_2_ treatment, but not all to the same extent (**Fig 3d**). Indeed, the relocalization of the C37S and C38S IL-38 mutants was remarkably decreased compared to the WT protein (**Fig 3e**).

Altogether, these data support a role for the C37-C38 disulfide bond in facilitating conformational changes and modification of the biochemical properties of IL-38 in response to oxidation.

### IL-38 forms granules in normal human epidermis

Finally, to determine the physiological relevance of our observations, we explored IL-38 aggregation in healthy human epidermis. Following immunostaining and confocal microscopy, 3D reconstruction of full-section stacks revealed the presence of granular IL-38 structures in epidermal keratinocytes (**Fig 4a**), strikingly resembling those observed in NHK/38 cells in oxidative conditions (**Fig 1b; Supp Fig 1c**). The specificity of IL-38 detection was confirmed, as described previously (Mermoud et al., 2022; Talabot-Ayer et al., 2019), by staining with an isotype control antibody (**Supp Fig 4**). The human epidermis is formed by keratinocytes arranged in 4 successive layers defined as basal, spinous, granular and cornified layers (**Fig 4b**). Each layer is characterized by specific morphological features reflecting the differentiation state of the keratinocytes, which increases from the basal to the cornified layer, where terminally differentiated cells undergo a specialized form of cell death, called cornification (Martin *et al.,* 2021). As previously described (Mermoud *et al.,* 2022), strong IL-38 staining was detected in all layers containing live cells, from the basal to the granular layer, while the signal was much lower in the cornified layer (**Fig 4a and Supp Fig 4**). Detection of IL-38 granules by automated image analysis (**Fig 4c**) and quantification of their fluorescence intensity (**Fig 4c; Fig 4d**) revealed an accumulation of granules with more intense fluorescence in the granular layer, which contains the live cells closest to the air interface.

**Figure 4.**
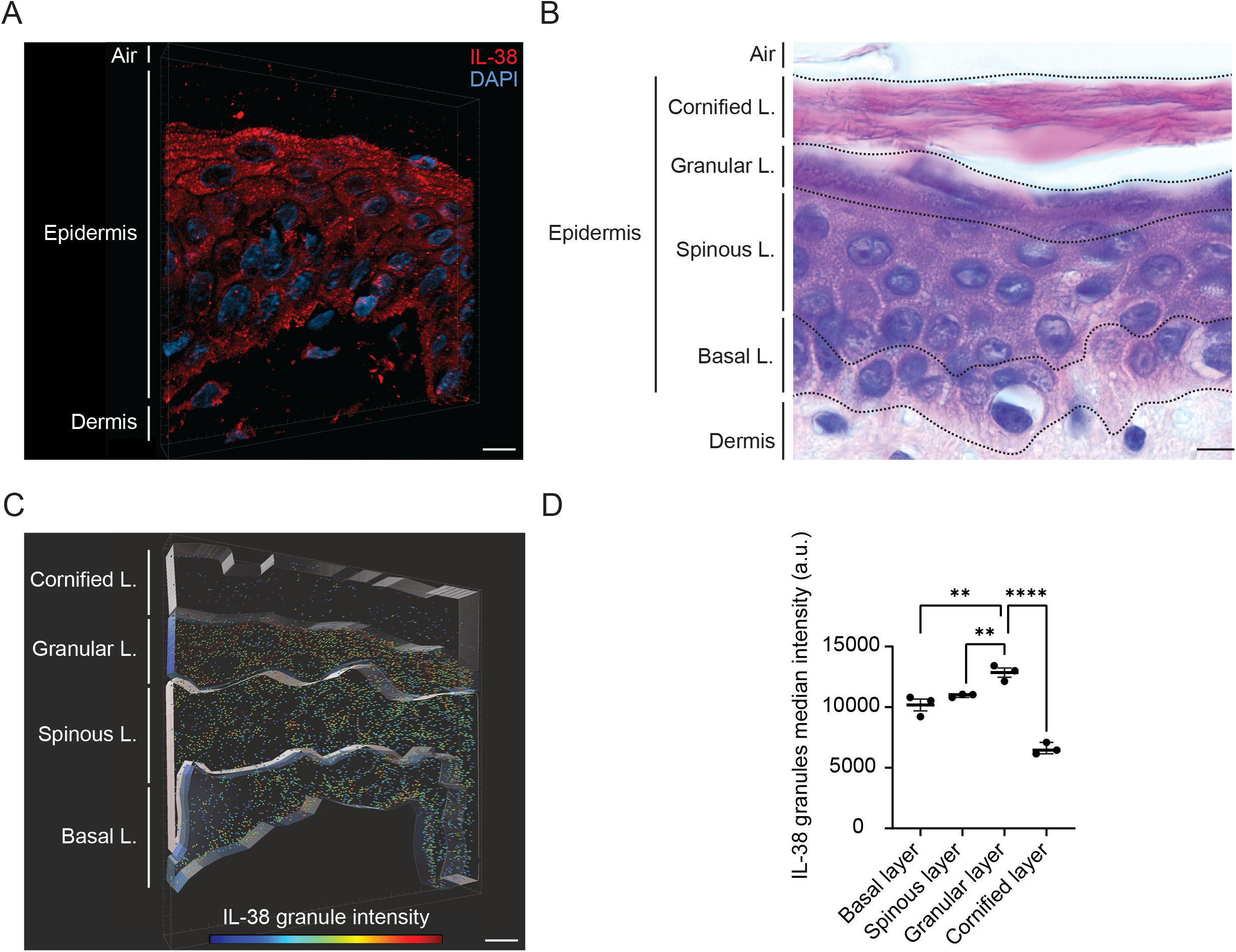
IL-38 forms granules in normal human epidermis. (A) Full-section reconstruction of a confocal microscopy stack of IL-38 immunostaining (red) in normal human epidermis. Dermis, epidermis and the epidermis-air interface are delimited on the left. DAPI (blue) was used to stain nuclei. The image is representative of n=3 different healthy donors. Original magnification 63x. Scale bar, 5 μm. (B) HE stained normal human skin section showing dermis, epidermis, and epidermal-air interface (indicated on the left). Dashed lines delimit the boundaries of the basal, spinous, granular and cornified layers. Original magnification 100x. Scale bar, 10 μm. L., layer (C) Detection of IL-38 granules in normal human epidermis using the “spots” module of Imaris. The color of the granules indicates their median fluorescence intensity in a range of 4008 (blue) to 20000 (red) arbitrary units. Epidermal layers are highlighted on the left and were defined manually with Imaris based on conventional morphological criteria. Scale bar, 5 μm. L., layer. (D) The median fluorescence intensities of all IL-38 granules in a given layer are shown as individual values and means + SEM for n=3 independent healthy donors. **p<0.01, ****p< 0,0001, by two-way ANOVA with Sidak’s multiple comparisons test. a.u., arbitrary units.

These data thus demonstrate the presence of granular IL-38 structures in healthy human epidermis, with highest IL-38 staining intensity in the viable keratinocytes that are most exposed to environmentally induced OS.

## Discussion

In this study, we describe oxidation-induced formation of amyloid aggregates as an intrinsic biological property of IL-38. Computational analysis revealed the presence of an LCR, including two ACs, in the human IL-38 protein sequence. LCRs and ACs are known to mediate the assembly of proteins into biomolecular condensates (Cereghetti *et al.*, 2018) and we indeed observed IL-38 relocalization to insoluble granular structures in human NHK keratinocytes during menadione-induced OS. As described also by others (Aulas et al., 2018), menadione treatment led to SG formation in NHK cells. In line with the previously reported association of IL-33 with SGs (Chen *et al.,* 2022), IL-38 condensates partially co-localized with G3BP1 positive structures. This observation is consistent with our previous identification of destrin, peroxiredoxin 1 and peroxiredoxin 2, which were detected in stress granules (Gwon et al., 2021; Jain et al., 2016), as IL-38 interaction partners (Talabot-Ayer *et al.*, 2019). Interestingly, IL-38 further localized to additional, G3BP1 negative cytoplasmic granules of currently unknown nature and function. Biomolecular condensates have emerged as key regulators of a myriad of cellular processes, such as signal transduction (Case et al., 2019; Du and Chen, 2018; Su et al., 2016), cell fate decisions (Samir et al., 2019), or the regulation of intracellular local protein concentration (Klosin et al., 2020). Similarly, the existence of functional amyloids, which have been primarily studied in microorganisms (Levkovich et al., 2021), is now increasingly recognized also in mammals, where they are involved in processes including skin pigmentation (Fowler et al., 2006), storage and secretion of peptide hormones (Maji et al., 2009), and intracellular signalling (Li et al., 2012). The role of IL-38 containing macromolecular complexes, or of IL-38 amyloids as such, in keratinocyte biology, as well as a potential contribution of other LCR-containing IL-1 family members thus clearly warrant further investigation.

Post-translational modifications, such as phosphorylation or acetylation, are known to regulate LCR-mediated protein aggregation (Cereghetti *et al.,* 2018; Saito et al., 2019). Interestingly, in the case of IL-38, amyloid formation was promoted by the oxidation of specific cysteines and our results are consistent with a model in which aggregation is favoured by the formation of an intramolecular disulfide bond. While disulfide bond formation was previously reported to enhance amyloidogenesis of different proteins, this was linked to the formation of inter-molecular disulfide bonds, which increased the molecular interactions between polypeptides. In contrast, intramolecular disulfide bond formation is generally believed to stabilize protein monomers and decrease amyloid formation (Mitra and Sarkar, 2022; van Dam and Dansen, 2020). To our knowledge, our current findings thus represent a first example of amyloid formation promoted by a destabilizing intramolecular disulfide bond. Creation of this bond between two vicinal cysteines is predicted to lead to a major conformational change that exposes aggregation-prone regions of the protein, thus representing a new mechanism for the regulation of amyloidogenesis through a redox switch.

Remarkably, in the epidermis of healthy human skin, IL-38 forms granular structures that closely resemble those detected in menadione-treated NHK/38 cells. A simple explanation would be that exposition of the epidermis to environmental OS promotes formation of amyloid IL-38 aggregates and higher-order granular structures. Interestingly, IL-38 has been linked to keratinocyte differentiation and epidermal cornification (Lachner *et al.,* 2017; Mermoud *et al.,* 2022). In this specialized form of programmed cell death, the nuclei of terminally differentiated keratinocytes are destroyed, while mRNA must remain available to allow the translation of proteins required for the assembly of the cornified envelope, which confers the main barrier function to the skin. Ribonucleoprotein (RNP) granules were proposed to serve as sinks for key mRNAs during keratinocyte enucleation, therefore acting as translation gatekeepers (Wotherspoon et al., 2020). Our current observation of IL-38 granules partially co-localizing with the typical RNP component G3BP1 (Yang et al., 2020) links IL-38 to RNP condensates and suggests that the formation of IL-38 aggregates could contribute to mRNA protection during cornification.

Overall, our results support a model in which oxidation-sensitive cysteines act as redox switches to modify the conformation of the IL-38 protein, and thus the surface exposure of its ACs, shuttling it from a soluble state into biomolecular condensates. The observation of IL-38 granules in human epidermal keratinocytes highly exposed to environmental OS further suggests that this previously unrecognized biochemical feature of IL-38 may be physiologically relevant at this epithelial barrier.

## Experimental procedures

Detailed methods are available in the Supplemental experimental procedures

### Cell culture

The Normal Human Keratinocyte cell line (Steenbergen et al., 1996) inducibly expressing human IL-38 (NHK/38 cells) (Talabot-Ayer et al., 2019) was grown in keratinocyte-serum free medium (K-SFM, Thermo Fisher Scientific AG, Waltham, MA, USA), supplemented with human recombinant epidermal growth factor and bovine pituitary extract (Thermo Fisher Scientific AG), 1% penicillin and streptomycin. HEK293T cells were cultured in Dulbecco’s Modified Eagle’s Medium (DMEM, 4.5 g/l glucose, Thermo Fisher Scientific AG, Waltham, MA, USA), supplemented with 10% FCS, L-glutamine and 1% penicillin and streptomycin, and transfected with expression vectors for WT, C37S and C38S IL-38.

### Immunofluorescence

NHK/38 cells and sections of normal human epidermis were stained with a monoclonal anti-hIL-38 antibody (H127C) (14-7385-82, Thermo Fisher Scientific AG, 1/2000), followed by an Alexa Fluor 594-labeled goat anti-mouse IgG2b (115-585-207, Jackson Immuno Research Europe Ltd, 1/200). Slides were imaged with a LSM800 confocal microscope (Carl Zeiss Microscopy, Feldbach, Switzerland) and the ZEN black software (Carl Zeiss Microscopy).

### All-atom accelerated molecular dynamics

IL-38 structural dynamics were studied through accelerated molecular dynamics (aMD) as implemented in Amber20 (Case et al., 2021). 1-microsecond production runs were simulated for fully reduced IL-38, C37-C38 DSB IL-38 and C37-C38+C2-C43 DSB IL-38.

## Supporting information

Supplemental data and experimental procedures

## Data availability

All data are available within the paper and its Supplementary information. Biological materials are available from the authors upon request.

## Author contributions

Conceptualization: A.D.B, G.C., Y.C., M.P., A.F.M., G.P. Formal analysis: A.D.B, G.C., S.K.J., Y.C., A.F.M. Funding acquisition: Y.C., M.P., A.F.M., G.P. Investigation: A.D.B., G.C., J.T., D.T.A., S.K.J., A.H., Y.C., A.F.M. Methodology: A.D.B, G.C., J.T., D.T.A., S.K.J., C.S., Y.C., M.P., A.F.M., G.P. Project administration: A.D.B., Y.C., M.P., G.P. Resources: C.S. Supervision: Y.C., M.P., G.P. Validation: A.D.B., G.C., S.K.J., Y.C., A.F.M. Visualization: A.D.B., G.C., S.K.J., Y.C., A.F.M. Writing – original draft: A.D.B., G.P. Writing – review & editing: A.D.B, G.C., A.H., C.S., Y.C., M.P., A.F.M., G.P. All authors reviewed the final manuscript and approved the submitted version.

## Acknowledgements

We would like to thank Ali Modarressi (University Hospitals, Geneva, Switzerland) for providing healthy human skin samples used in this study. We would like to thank Sergei Startchik and Nicolas Liaudet (Bioimaging Core Facility, Faculty of Medicine, University of Geneva, Switzerland) for their help with microscopy image analysis. We would like to thank the Protein Platform (Faculty of Medicine, University of Geneva, Switzerland) for their help with the production of recombinant IL-38 proteins and the Proteomics Core Facility (Faculty of Medicine, University of Geneva, Switzerland) for their help with the analysis of N-terminal IL-38 peptides. This work was supported by grants from the Swiss National Science Foundation (310030_188470 to GP), the Rheumasearch Foundation, the Kurt and Senta Herrmann Foundation, the Medicor Foundation, and by a generous donor advised by Carigest SA. Work in the Peter-laboratory is supported by funding from the Swiss National Science Foundation (SNSF), the Synapsis Foundation Switzerland, and the Department of Biology of ETH Zürich. Antonio Francés-Monerris was supported by the grant PID2021-127554NA-I00 funded by MCIN/AEI/10.13039/501100011033 and by “ERDF A way of making Europe”. The proteomic experiments were partially supported by the Agence Nationale de la Recherche under projects ProFI (Proteomics French Infrastructure, ANR-10-INBS-08) and GRAL, a program from the Chemistry Biology Health (CBH) Graduate School of University Grenoble Alpes (ANR-17-EURE-0003).

