## Supplemental data and experimental procedures for "Oxidation-sensitive cysteines drive IL-38 amyloid formation"

#### Supplemental experimental procedures

##### LCR and AC determination

The amino acid (aa) sequences of IL-1 family members were retrieved from the Uniprot Database (https://www.uniprot.org/) using the following identifiers: IL-1 beta, P01584; IL-1 alpha, P01583; IL-33, O95760; IL-36 alpha, Q9UHA7; IL-36 beta, Q9NZH7; IL-36 gamma, Q9NZH8; IL-18, Q14116; IL-1 receptor antagonist, P18510; IL-36 receptor antagonist, Q9UBH0; IL-37, Q9NZH6; and IL-38, Q8WWZ1. Sequences were analyzed using the SEG algorithm (ref. Wootton et al. 1994) to identify LCRs. The predictions were run using the following parameters: trigger window length [W] = 25, trigger complexity [K(1)] = 3.0 and extension complexity [K(2)]) = 3.3, or [W] = 12, [K(1)] = 2.2, and [K(2)] = 2.5. The aa sequence of IL-38 was further analyzed using the amyloid-predicting consensus tool AmylPred2.0 (Tsolis et al., 2013). AmylPred2.0 compiles the prediction of 11 different methods developed to identify amyloidogenic regions. A region was scored as an AC if a consensus of five or more methods was reached.

##### Plasmid mutagenesis

Plasmid pcDNA4/TO/hIL-38 (Talabot-Ayer et al., 2019) was used as a template to generate the plasmid pcDNA4/TO/hIL-38-TD, encoding the major T44 and D51 IL-38 variant present in the European population (Ensembl entry: ENSG00000136697), by PCR-directed mutagenesis as previously described (Georg et al., 2020). In turn, pcDNA4/TO/hIL-38-TD was used as a template to generate the C37S (pcDNA4/TO/hIL-38-TD-37S) and C38S (pcDNA4/TO/hIL-38-TD-38S) IL-38 mutants.

##### Cell culture, transfection and treatment

The immortalized Normal Human Keratinocyte cell line (Steenbergen et al., 1996) inducibly expressing human IL-38 (NHK/38 cells) (Talabot-Ayer *et al.*, 2019), was grown in keratinocyte-serum free medium (K-SFM, Thermo Fisher Scientific AG, Waltham, MA, USA), supplemented with human recombinant epidermal growth factor and bovine pituitary extract (Thermo Fisher Scientific AG), 1% penicillin and streptomycin, at 37˚C in a humidified atmosphere containing 5% CO2. NHK/38 cells were plated in 6-multiwell plates at a density of $7x10^5$ cells/well and IL-38 expression was induced with doxycycline (Dox) at 1 µg/ml for 24h. A model of oxidative stress (OS) was established by treating Dox-induced NHK/38 with the oxidant menadione (Aulas et al., 2018) (Merck, Darmstadt, Germany) dissolved in 99% ethanol at 75 µM for the indicated times. Chloroquine (Merck, Darmstadt, Germany) was diluted in KSFM medium and added at 250 µM for 24 h to Dox-treated NHK/38 cells to induce autophagy (Mauthe et al., 2018). Staurosporine (Merck, Darmstadt, Germany) was diluted in DMSO and added at 1 µM for 5 h to Dox-treated NHK/38 cells to induce apoptosis (Jacobson et al., 1993).

Human embryonic kidney (HEK) 293T cells were cultured in Dulbecco′s Modified Eagle′s Medium (DMEM, 4.5 g/l glucose), supplemented with 10% FCS, L-glutamine, 1% streptomycin and penicillin. HEK293T cells were seeded in 6 well plates at a density of $3x10^5$ cells/well and transiently transfected 24 hours later, at a confluence of 70-80 %, with pcDNA4/TO/hIL-38-TD, pcDNA4/TO/hIL-38-TD-37S or pcDNA4/TO/hIL-38-TD-38S, using the JetPrime reagent (Polyplus, Illkirch, France) following the manufacturer's instructions. To induce OS, HEK293T cells were treated with 10 mM $H_2O_2$ for 1 h.

**Human skin samples**

Skin biopsies were taken from healthy adults undergoing surgery at the Department of Plastic and Reconstructive Surgery of the Geneva University Hospitals in Switzerland. This study was conducted according to the Declaration of Helsinki, and approved by the local ethics committee of the University Hospitals of Geneva (protocol numbers 2017-00700, 2020-01102). Written informed consent was obtained from each individual. Skin tissue samples were fixed in 4%

formaldehyde and embedded in paraffin. Five µm sections were cut and stained with hematoxylin and eosin (HE) or processed for immunofluorescence.

**Immunofluorescence**

NHK/38 cells were seeded in chamber slides at a density of $3.5x10^4$ cells/well and incubated after 24 h with 1 µg/ml Dox for one additional day, before treatment with 75 µM menadione, 250 µM chloroquine or 1 µM staurosporine for indicated times. Cells were fixed (PBS, 3.7% paraformaldehyde), permeabilized (PBS, 0.1% Triton X-100) and washed (Dako wash buffer, Agilent Technologies AG), before staining. For extraction of the soluble cellular content, before fixation as described above, cells were incubated for 1 minute with extraction solution (100 mM PIPES, pH 6.9; 1 mM $MgCl_2$; 1 mM EGTA; 1 % Triton X-100; 30% glycerol) supplemented with 2 µM phalloidin (Merck, Darmstadt, Germany) and washed two times with PEM buffer (100 mM PIPES, pH 6.9; 1 mM $MgCl_2$; 1 mM EGTA) supplemented with 2 µM phalloidin. The antibodies used for immunostaining were: a monoclonal mouse IgG2b, kappa anti-hIL-38 antibody (H127C) (14-7385-82, Thermo Fisher Scientific AG, 1/2000), polyclonal rabbit anti-G3BP1 (PA5-29455, Thermo Fisher Scientific AG, 1/1000), monoclonal rabbit [EPR4844] anti-p62 (ab109012, Abcam, Cambridge, UK, 1/400) and polyclonal rabbit anti-cleaved caspase-3 (D175) (9661S, Cell Signaling Technology, MA, USA, 1/400) in antibody diluent (Dako, Agilent Technologies AG) overnight at 4˚C. Subsequently, primary antibodies were detected with Alexa Fluor 594-labeled goat anti-mouse IgG2b (115-585-207, Jackson Immuno Research Europe Ltd, 1/200), and/or Alexa Fluor 488-labeled donkey anti-rabbit IgG (711-545-152, Jackson Immuno Research Europe Ltd, 1/200), as appropriate, and slides were stained with 4',6-diamidino-2-phenylindole (DAPI; D3571,Thermo Fisher Scientific AG, 1/ 2000 in PBS) and mounted in FluoreGuard medium (ScyTek Laboratories, Inc., Logan, UT). To assess staining specificity, negative controls were performed in absence of primary antibodies or with isotype-matched control antibodies. In addition, the specificity of IL-38 detection was previously assessed by using control NHK cells lacking IL-38 expression (Talabot-Ayer *et al.*, 2019). Slides were imaged with a LSM800 confocal microscope (Carl Zeiss Microscopy, Feldbach, Switzerland) and the ZEN black software (Carl Zeiss Microscopy).

Human skin biopsies were processed as described (Talabot-Ayer *et al.*, 2019). The antibody used for IL-38 staining was: monoclonal mouse IgG2b, kappa anti-hIL-38 antibody (H127C) (14-7385-82, Thermo Fisher Scientific AG, 1/2000), followed by Alexa Fluor 594-labeled goat anti-mouse IgG2b (115-585-207, Jackson Immuno Research Europe Ltd, 1/200). DAPI was used to stain the nuclei and full-section stacks were acquired with a resolution of 0.5 µm in the xy plane and 0.15 µm in the z axis. Specificity of the staining was evaluated by comparison with an isotype matched IgG control in the same skin sections. Slides were imaged with a LSM800 confocal microscope (Carl Zeiss Microscopy, Feldbach, Switzerland) and the ZEN black software (Carl Zeiss Microscopy). IL-38 granule detection and classification, as well as epidermal layer definition in 3D stacks of healthy human skin was done with the image processing software Imaris. Epidermal layers were defined by conventional morphological criteria (Gilbert et al., 2000) using the transmitted light detection and DAPI channels. The basal layer was defined as the single line of cells in contact with the epidermal-dermal interface. The spinous layer was defined to include polyhedral cells located between the boundaries of the basal and granular layers, and by the presence of a high number of desmosomes between cells. The granular layer was defined by the transition from polyhedral to flat cells and the decrease in the number of desmosomes. The cornified layer was defined based on the loss of nuclei and normal cell morphology. Automatic granule detection by the Imaris spots module was supervised by a specialized researcher and the quality threshold was set at 1600 for the 3 donors. For graphical representation, granules were colored based on IL-38 median fluorescence intensity within each granule, in a range of 4008 to 20000 arbitrary units in all epidermal layers.

**Cell fractions and Western blotting**

Subcellular fractionation was performed with a Subcellular Protein Fractionation Kit for Cultured Cells (Thermo Fisher Scientific AG, Waltham, MA, USA). Manufacturer's instructions were followed except for the following variations: all solutions of the kit, as well as the PBS used for washing, were supplemented with 40 mM iodoacetamide to block free cysteines and thus avoid artifactual aggregation. An additional washing step as compared to recommendations was introduced between isolation of each fraction. The insoluble fraction was defined as the fraction remaining after extraction of the cytoplasmic, membrane, nuclear, chromatin bound and cytoskeletal fractions, and it was denatured with 200 µl denaturing urea buffer (30 mM Tris-HCl, 7 M urea, 2 M thiourea, 4% CHAPS, 40 mM iodoacetamide, 2% β-mercaptoethanol (β-ME)) by incubating for 10 min at 37 ºC and vortexing every 2 min at maximum power for 10-15 s, prior to centrifugation for 10 min at 4°C and 20,000 g for clearing. 10% of the total volume of cytoplasmic and insoluble fractions was diluted in reducing Laemmli buffer and separated electrophoretically on a 4-20% gradient Bis-Tris SDS-PAGE (mPAGE; Merck KGaA, Darmstadt, Germany) with MES running buffer, and blotted onto PVDF membranes. The membranes were blocked with 5% horse serum in TBS, 0.05% Triton X-100 (TBST) and probed with biotinylated polyclonal goat anti-IL38 antibody (BAF2427, R&D Systems, 1/1000). Immunoreactive bands were detected using appropriate HRP-labeled secondary reagents for Clarity Western ECL detection (Bio-Rad Laboratories, Inc., Hercules, CA) on a LAS4000 imager (Fujifilm Life Science, Düsseldorf, Germany). Stripping was carried out with Re-Blot Plus Strong antibody stripping solution (Merck & Cie) for 20-40 min at RT and blocking with TBST, 5% horse serum before reprobing with a monoclonal mouse α-

Tubulin antibody (sc-23948, Santa Cruz, TX, USA; 1/500), in TBST. Western blot (WB) densitometry analysis was carried out with the ImageJ software (Schneider et al., 2012).

**LDH viability assay**

NHK/38 cells were plated in 48-multiwell plates at a density of $1.8x10^5$ cells/well and incubated with 1 µg/ml Dox on the following day for 24 h. Cells were then treated with 75 µM menadione or ethanol, used as a control, for 2 h or 24 h, before supernatants were harvested and tested with the Cytotoxicity Detection Kit (LDH) (Merck KGaA, Darmstadt, Germany) following the manufacturer's instructions. As positive control for cell death, one well of control NHK/38 cells was lysed with 1% Triton X-100 in K-SFM medium.

**Recombinant protein expression and purification**

For the production of the recombinant C-terminally His-tagged human WT IL-38 aa1-152 used to study amyloid formation, the complete human IL-38 cDNA sequence was codon optimized for *E. Coli.*, and synthetized and cloned by GenScript Biotech (Piscataway, NJ, USA) into a modified pET30-a (+) vector (Addgene, Watertown, MA, USA) in order to add a C-terminal 6xHis tag. The protein was expressed in BL21 (DE3) *E. Coli.* cells grown in TB media (Formedium, Swaffham, England) until reaching a $OD_{600}$ of 0.5-0.7 after which protein expression was induced by adding 500 µM IPTG (Euromedex, Souffelweyersheim, France) and grown O.N at 22°C. Pelleted cells were resuspended in a strongly reducing lysis buffer (50 mM Tris pH 8, 500 mM NaCl, 10 mM β-ME, 0.5% Triton X100) + Complete protease inhibitor (Roche, Basel, Switzerland) before being disrupted by pulsed sonication for a total of 6 minutes (30 sec on, 30 sec off cycles at 50% amplitude). After lysis, 5% glycerol was added before a 1h centrifugation at 4°C. The harvested supernatant was batch incubated for 30 min with 7ml Ni-NTA resin (Qiagen, Germantown, MD, USA), washed 4*40 ml with lysis buffer supplemented with 20 mM imidazole and eluted in 50 mM Tris pH 8, 500 mM NaCl, 2 mM β-ME, 5% glycerol and 200 mM imidazole. Recovered fractions were pooled and dialysed O.N using a 10 kDa dialysis cassette (Thermo Fisher Scientific AG, Waltham, MA, USA) against 50 mM Hepes pH 7, 50 mM NaCl, 5 % glycerol and 2 mM β-ME (Mono-Q buffer1). Following dialysis, the sample was directly loaded on a 5 ml HiTrap Q HP column (GE healthcare, Chicago, IL, USA), washed for 30 ml with buffer1 then eluted using a 50 ml gradient bringing the NaCl concentration to 1.5 M. Fractions of interest were pooled based on SDS–polyacrylamide gel electrophoresis (PAGE) analysis, concentrated in a 10 kDa Amicon concentrator (UFC901024, Sigma-Aldrich, MO, USA) and injected on a S75 10/300 GL increase column (GE Healthcare, Chicago, IL, USA) running in 30 mM Hepes pH 7, 500 mM NaCl, 1 mM β-ME and 5% glycerol. Following a final PAGE quality control, purified IL38 was concentrated to 1 mg/ml and sterile filtered using a 0.2 μm PES filter and frozen at -80°C.

Bottom-up nanoliquid chromatography-tandem mass spectrometry (nanoLC-MS/MS) performed on in-gel Lys-C digested IL-38 immunoprecipitated from NHK/38 cells revealed N-terminal IL-38 peptides starting at cysteine (C)2, in agreement with previous observations in apoptotic A549 lung carcinoma cells (Mora et al., 2016). N-terminal methionine excision, which likely accounts for the generation of this aa2-152 form of IL-38, is a common post-translational modification in eukaryotic cells, suggesting that aa2-152 IL-38 corresponds to a predominant naturally occurring form of the protein. To produce recombinant human WT IL-38 aa2-152 used for disulfide mapping, the human aa2-152 IL-38 cDNA sequence was codon optimized for *E. Coli*, synthetized and cloned by GenScript Biotech into a pET21a vector (Addgene, Watertown, MA, USA) in order to add Tobacco Etch Virus (TEV) protease cleavable N-terminal 6xHis and Strep-tag II tags. The protein was expressed in Rosetta (DE3) *E. Coli* cells (70954, Sigma-Aldrich, MO, USA). For purification, cell pellets were resuspended in lysis buffer (50 mM Tris (pH 8.0), 150 mM NaCl, and 3 mM β-ME) supplemented with an anti-protease cocktail (Complete EDTA-free, Roche, Basel, Switzerland). Lysis was performed with a Microfluidizer LM-20 set at 18’000 psi, and lysates clarified by centrifugation. Affinity purification was carried out on a Strep-Tactin XT 4Flow column (2-5028-001, IBA Lifesciences GmbH, Göttingen, Germany) and in-house-purified Tobacco Etch Virus (TEV) protease was added for on-column digestion of the N-terminal tag. Next, a His-trap FF column (17525501, Cytiva, MA, USA) was plugged after the STREP-Tactin column and the cleaved IL-38 protein was eluted in lysis buffer supplemented with 10 mM imidazole. Finally, the eluate was concentrated using a 10 kDa MW cutoff Amicon concentrator (UFC901024, Sigma-Aldrich, MO, USA) and size exclusion chromatography (SEC) was carried out at 4°C on a Superdex 200 10/300 column (28990944, Cytiva, MA, USA). Fractions were analyzed by SDS-PAGE and fractions containing pure IL-38 protein were flash-frozen in liquid nitrogen and stored at -80 °C.

For the production of WT and C38S IL-38 aa2-152 proteins used to study amyloid formation, the purification strategy was adapted in order to increase the yield. The full-length WT or mutant human IL-38 cDNA sequence was codon optimized for *E. Coli*, synthetized and cloned by GenScript Biotech into a pET21a vector (Addgene, Watertown, MA, USA) in order to add a TEV protease cleavable N-terminal 6xHis tag. Proteins were expressed in LOBSTR BL21 (DE3) *E. Coli* cells (EC1002, Kerafast, MA, USA) and purified under reducing conditions (7.5 mM β-ME, except for dialysis at 15 mM β-ME). Pellets were resuspended in lysis buffer (50 mM Tris pH 8.0, 500 mM NaCl, 10 mM imidazole, 5% glycerol, 0.5% Triton X-100) supplemented with an anti-protease cocktail. Lysis was performed with a Microfluidizer LM-20 set at 18'000 psi and lysates clarified by centrifugation. Affinity purification was carried out on a His-trap FF column (17525501, Cytiva, MA, USA) and the N-terminal tag was cleaved with His-TEV protease during dialysis against 20 mM Tris pH 8; 250 mM NaCl; 5% glycerol. Then, proteins of interest were separated from contaminants (uncleaved protein, TEV protease, tag) on a His-trap FF column, and purified on a Q-HP column (17115401, Cytiva, MA, USA). Fractions were analyzed by SDS-PAGE, IL-38 containing fractions were pooled, concentrated on 10 kDa MW cutoff Amicon concentrators (UFC901024, Sigma-Aldrich, MO, USA) and SEC was carried out on a Superdex 75, 10/300 column (29148721, Cytiva, MA, USA) equilibrated in sterile SEC buffer (30mM Hepes pH 7, 500mM NaCl, 5% glycerol, 1 mM β-ME). Fractions were analyzed by SDS-PAGE and fractions containing pure proteins were pooled, flash-frozen in liquid nitrogen and stored at -80 °C.

Cdc19 was purified as previously described (Cereghetti et al., 2022b). In short, *E. Coli* cells were transformed with a plasmid expressing wild-type Cdc19-strep and grown at 37 °C in LB media (1% peptone, 0.5% yeast extract, 0.5% NaCl) containing 30 µg/ml chloramphenicol and 100 µg/ml carbenicillin. When $OD_{600}$ 0.6 was reached, protein expression was induced by addition of IPTG to a final concentration of 0.1 mM. Cells were grown at 16 °C for 12 h, harvested by centrifugation. Then, they were resuspended in cold purification buffer (100 mM Tris-HCl pH 7.4, 200 mM NaCl, 1 mM MgCl2, 10% glycerol, 1 mM phenylmethylsulfonyl fluoride (PMSF), 1 mM DTT) supplemented with protease inhibitor tablets (Roche, Basel, Switzerland) and 75 U/ml of Pierce universal nuclease (Thermo Fisher Scientific, Waltham, MA, USA), and lysed by freezer milling (SPEX SamplePrep 6870 Freezer/Mill). Subsequently, lysates were cleared by centrifugation (4 °C, 30 min, 48000 g), and the supernatant was loaded on a Strep-Tactin Superflow Plus column (Qiagen, Germantown, MD, USA) at 4 °C following the manufacturer's instructions. Proteins were eluted using purification buffer supplemented with 2.5 mM desthiobiotin, and aliquots were stored at -80 °C.

**Congo red and Thioflavin T staining**

Staining with Congo Red (CR) and Thioflavin T (ThT) was performed essentially as previously described (Cereghetti et al., 2022a; Cereghetti et al., 2022b). Congo red (75768-25MG, Sigma-Aldrich, MO, USA) or Thioflavin T (T3516, Sigma-Aldrich, MO, USA) were dissolved in water to a final concentration of 1 mM or 2.5 mM, respectively, and filtered (0.2 µm; Millipore). Purified proteins were thawed on ice, cleared by centrifugation (10 min, 21,000 g,

4 °C), and diluted to 10 μM in purification buffer (30mM Hepes pH 7, 500mM NaCl, 5% glycerol, 1 mM β-ME). $H_2O_2$ (Sigma-Aldrich, MO, USA) was added as indicated in the figures. After addition of the CR or ThT solution (1:10 dilution), fluorescence intensity was recorded for up to 18 h at 37 ºC in a 384-well plate (Corning Life Sciences, Corning, NY USA) using a CLARIOstar plate reader (BMG Labtech, Ortenberg, Germany). For CR, the fluorescence intensity was measured at 614 nm with an excitation at 560 nm. For ThT, excitation was set at 450 nm and emission spectra were recorded at 490 nm.

**SDD-AGE assay**

HEK293T cells transfected with pcDNA4/TO/hIL-38-TD were treated 24 h after transfection with 10 mM $H_2O_2$ for 1 h, or left untreated. Before harvesting, cells were washed with 40 mM iodoacetamide in PBS to block free sulfhydryls and avoid artifactual aggregation during lysis. SDD-AGE analysis was performed essentially as previously described (Cereghetti *et al.*, 2022a). Briefly, pellets were resuspended in 1:1 volume of lysis buffer (50 mM Tris, pH 7.5; 150 mM NaCl; 1% (vol/vol) TritonX-100; 2.5 mM EDTA; 0.33 mM PMSF; 20 mM iodoacetamide; protease inhibitor cocktail) and mechanically disrupted with metal beads (30 Hz, 3 min) at RT. Next, cell debris were removed by mild centrifugation (2000 rpm, 2 min, 4 °C), and supernatants were mixed with 3 volumes of sample buffer (2X TBE, 20% glycerol, 8% SDS, bromophenol blue). Electrophoresis was performed in a 1.5% agarose TBE gel supplemented with 0.1% SDS (running buffer: 1X TBE, 0.1 % SDS). Proteins were transferred onto a nitrocellulose membrane by capillarity overnight, followed by blocking and blotting for IL-38 as described above.

**Top-down nanoliquid chromatography coupled to tandem mass spectrometry analyzes**

Purified recombinant IL-38 protein was alkylated or not using a 30 min incubation in the dark with 50mM iodoacetamide in 25mM ammonium bicarbonate. Solutions containing 1 µM of alkylated or non-alkylated forms were diluted 10 times in 5% acetonitrile, 0.1% trifluoroacetic acid and analyzed by online nanoliquid chromatography coupled to tandem mass spectrometry (nanoLC-MS/MS) (Ultimate 3000 RSLCnano and Q-Exactive Plus, Thermo Fisher Scientific AG, Waltham, MA, USA). For this purpose, the proteins were sampled on a precolumn (300 nm x 5 mm C4 PepMap, Thermo Fisher Scientific, Waltham, MA, USA) and separated in a 75 µm x 250 mm C4 column (ReproSil-Pur 300, 3µm, Dr. Maisch) using a 20 min gradient at a flow rate of 300 nl/min. MS and MS/MS data were acquired using Xcalibur version 3.1 (Thermo Fisher Scientific, Waltham, MA, USA). The non-alkylated and alkylated forms of IL-38 were analyzed in mass ranges of respectively 600 to 1600 m/z and 600 to 2000 m/z, with MS1 resolutions of respectively 140’000 and 70’000 (at m/z 200), and with MS2 resolution of 70’000 (at m/z 200). Parallel reaction monitoring experiments were conducted to analyze two major charge states observed in MS1 spectra, respectively m/z 1123.56 and 1532.06 corresponding to 16+ and 9+ forms for non-alkylated IL-38, and m/z 1701.9 and 1891.04 corresponding to 10+ and 9+ forms for alkylated IL-38. All considered experimental masses were monoisotopic and non-protonated masses. The spectra were deconvoluted using the Xtract module in Freestyle (Thermo Fisher Scientific, Waltham, MA, USA). ProSightLite v1.4 (Northwestern University) was then used to match the experimental MS/MS fragments to the theoretical sequence of IL-38 (possible modifications: proton loss and carbamidomethylation on C residues).

**All-atom accelerated molecular dynamics simulations and molecular graphics**

*Preparation of the systems and general settings*

The IL-38 protein was prepared from the PDB ID: 5BOW. The initiator methionine 1 was removed following our and previous observations (Mora *et al.*, 2016) suggesting that aa2-152 IL-38 corresponds to a predominant naturally occurring form of the protein (see the recombinant protein expression and purification section). The H++ online application (Anandakrishnan et al., 2012) was used to obtain the correct protonation state at pH = 7. The resulting PDB structure was subsequently solvated by using the AmberTools20 toolset (Case et al., 2021) into a cubic water box with a size of 66.461x64.744x65.734 Å, containing 6548 water molecules and five $K^+$ counterions to neutralize the solvated system. The total number of atoms is 21979.

The protein was described with the ff99SB (Hornak et al., 2006) force field and water molecules and $K^+$ ions were described with the TIP3P parameters (Jorgensen et al., 1983). The system was initially energy-minimized for 6000 cycles of the steepest descent algorithm, followed by another 6000 cycles of conjugate gradient minimization. Later, the system was slowly heated up keeping the volume constant (NVT ensemble) during 200 ps up to a reference temperature of 310.15 K. Accelerated molecular dynamics (aMD, see details below) production runs were conducted at a constant pressure (NPT ensemble) of 1 atm employing the Monte Carlo barostat, whereas temperature conservation at 310.15 K was ensured through Langevin dynamics. Simulations were performed under periodic boundary conditions and particle mesh-ewald (PME) summation with a cut-off of 9.0 Å. The total simulation time for each run was 1 μs, and the employed time step was 1 fs. The SHAKE algorithm (Ryckaert et al., 1977) was applied to freeze bond distances involving H atoms. Atomic coordinates and velocities were stored every 40 ps, making a total of 25000 snapshots per trajectory usable for analyzes. Several 1-μs aMD replicas were conducted departing from the same initial structure and randomizing initial velocities at 310.15 K. All simulations have been performed with a GPU running a cuda based version of the Amber20 software (Case et al., 2021; Gotz et al., 2012; Salomon-Ferrer et al., 2013).

The IL-38 protein containing one covalent disulfide bond (DSB) between C37 and C38 (C37-C38) was prepared by selecting an appropriate snapshot from the fully reduced IL-38 simulations with a small S-S interatomic distance between the sulfur atoms of interest. The hydrogen atoms of the involved –SH groups in the DSB were subsequently removed, and the covalent S-S bond defined. The resulting C37+C38 DBS IL-38 protein was re-solvated and simulated by means of the same protocol as described above for the fully reduced IL-38. Similarly, an adequate snapshot from a simulation of C37+C38 DSB IL-38 with a close distance between the S atoms of the aa Cys2 and Cys43 was selected to prepare the protein with two DSBs, namely between C37 and C38 and C2 and C43 (C37-C38 + C2-C42), respectively. The same protocol was followed to solvate, neutralize the system, and simulate the protein dynamics.

*S-S distances and solvent-accessible surface area monitoring*

Interatomic sulfur-sulfur distances of interest were measured over time making use of the cpptraj program distributed in Ambertools20 (Case et al., 2021). Time series of the solvent-accessible surface area (SASA) values were calculated for the two AC regions identified through computational analysis (aa sequences 98-103 and 119-123, respectively). The S-S distances and the SASA time series for each 1 μs aMD trajectory were then plotted as frequency histograms making use of the Gnuplot 5.2 program.

*Accelerated molecular dynamics details*

Production simulations were run making use of the aMD method (Hamelberg et al., 2007; Hamelberg et al., 2004; Pierce et al., 2012) as implemented in Amber20 (Case et al., 2021). aMD drastically increases the conformational sampling up to three orders of magnitude (Feher et al., 2019; Kamenik et al., 2018; Pierce et al., 2012) by reducing energy barriers separating different states of a system. aMD is a well-established enhanced sampling method used in the study of protein structural dynamics (Calvo-Tusell et al., 2022; Curado-Carballada et al., 2019; Dodd et al., 2018; Heidari et al., 2019; Viswanathan et al., 2020). aMD parameters were obtained from a 1 μs conventional MD trajectory of the fully reduced IL-38 protein with the ff99SB/TIP3P force field, as previously described (Pierce et al., 2012). The parameters, in kcal/mol, are E(tot) = -62199, Alpha(tot) = 2080, E(dih) = 2080, and Alpha(dih) = 106. Three replicas of 1 μs aMD simulations were run for the reduced IL-38 protein and three for C37-C38 DSB IL-38. Analysis of the obtained S-S distances and SASA values revealed very consistent results for each system. One 1 μs aMD simulation was conducted for the C37-C38 + C2-C42 DSB IL-38 protein.

*Molecular graphics*

Molecular graphics were performed with UCSF Chimera, developed by the Resource for Biocomputing, Visualization, and Informatics at the University of California, San Francisco, with support from NIH P41-GM103311.

**Statistical analyzes**

The following tests were used with GraphPad Prism version 8.0.0 for Windows (GraphPad Software, San Diego, California USA), as indicated in the figure legends: two-way ANOVA with Sidak's multiple comparisons test, unpaired Student's t-test and Friedman test with Dunn's multiple comparisons test. Statistical significance was defined at a p-value < 0.05.0p

#### Supplemental Figure Legends

##### Fig. S1. IL-38+ granules form independently of autophagy, apoptosis, stress granule assembly, or loss of plasma membrane integrity in response to menadione

(A) LDH activity was measured in supernatants of Dox-induced NHK/38 cells after 2 and 24 h treatment with 75 µM menadione (square symbols) or vehicle (triangular symbols). Results are expressed as mean percentage with respect to LDH activity in the whole cell lysate (WCL) of 2h control cells ± SEM for n=3 independent experiments. **p<0.01, by two-way ANOVA with Sidak's multiple comparisons test. (B) Localization of p62 (white staining) was examined by immunofluorescence (IF) in Dox-induced NHK/38 cells treated with 0.25 mM chloroquine (CHQ) or its vehicle (NT) for 24 h (upper panels) or with 75 µM menadione (Men) or its vehicle (NT) for 2 h (lower panels). Scale bars: 5 µm. Results are representative of n=2 independent experiments. (C) Localization of cleaved caspase-3 (white staining) was examined by IF in Dox-induced NHK/38 cells treated with 1 µM staurosporine or its vehicle (NT) for 5 h (upper panels) or with 75 µM menadione (Men) or its vehicle (NT) for 2 h (lower panels). DAPI was used to stain the nuclei (blue staining). Scale bars: 5 µm. Results are representative of n=2 independent experiments. (D) Localization of IL-38 (red staining; second and fourth columns) and G3BP1 (green staining; third and fourth columns) was examined by confocal IF microscopy in Dox-induced NHK/38 cells treated with 75 µM menadione (Men) or its vehicle (NT) for 1 h. The soluble content of the cell was left intact (NO EXTRACTION; upper panels) or extracted (EXTRACTION; lower panels). Phalloidin was used to stabilize the actin cytoskeleton in extracted cells. Overlap between the red and green fluorescence signals is visible in yellow in the merged images (fourth column). DAPI (blue staining; first column) was used to stain the nuclei. IL-38 partially colocalized with G3BP1 in non-extracted cells and IL-38 staining remained detectable in G3BP1+ stress granules (white, solid arrows), in granules staining for IL-38, but not for G3BP1 (dashed, open arrows), and in nuclei in extracted menadione-treated cells. Scale bars: 5 µm.

**Fig. S2. Oxidation specifically triggers IL-38, but not Cdc19 amyloid formation**

C-terminally His-tagged full-length recombinant human IL-38 protein (left panel) was exposed to $H_2O_2$ (0.01, 0.1, 1 and 10 mM, grey to black symbols), or left untreated (lightest grey), and the *in vitro* formation of IL-38 amyloids was assessed by ThT staining and fluorescence detection during 13 h at 37 ºC. ThT alone (right panel) was exposed to $H_2O_2$ (0.01, 1 and 10 mM, grey to black symbols), or left untreated (lightest grey) during 13 h at 37 ºC, in order to exclude unspecific detection of ThT fluorescence. Results in both panels are expressed as mean ± SEM for n=3 independent experiments. A.u., arbitrary units. ThT, thioflavin T.

**Fig. S3. IL-38 disulfide bridge prediction, mapping and link to AC surface exposure**

(A) Histogram of the most relevant sulfur-sulfur (S-S) distances (X axis, in Å) and their frequency (Y axis), measured along a 1 μs accelerated molecular dynamics (aMD) simulation based on the human IL-38 crystal structure (PDB ID: 5BOW). The black dotted line indicates a distance of 4.5 Å, at which no solvent molecule can be placed between the two sulfur atoms and which can thus be considered as a hotspot for disulfide bridge formation. Results are representative of n=3 independent simulations (B) Top-down nanoLC-MS analyzes of purified recombinant human IL-38 aa2-152 submitted (bottom panel), or not (top panel), to alkylation. Linear IL-38 has a theoretical mass of 16833.23 Da. The observed mass (16829.17 Da) before alkylation (top panel) suggests the presence of 2 disulfide bridges (- 4 Da). The observed mass (17000.26 Da) after alkylation (bottom panel) indicates the presence of 2 disulfide bridges (-4Da) and 3 alkylated residues (+ 171Da). (C) Alkylated IL-38 was submitted to nanoLC-MS/MS analysis. The fragmentation pattern (b ions shown in blue and y ions shown in red) strongly suggests that C67, C70 and C123 are alkylated (open squares), whereas 2 disulfide bridges (open circles) occur in the N-terminal part of the protein, involving C2, C37, C38 and

C43. (D) Ribbon diagrams (left images) and surface representation (right images) from a 1 µs aMD simulation of disulfide-bonded (DSB) IL-38 carrying two disulfide bonds (C37-C38 and C2-C43). The two ACs are shown in green and cysteine residues in yellow. Disulfide bonds are indicated by black arrows. DSB, disulfide bonded. (E) Quantification of the surface exposition, represented by the solvent-accessible surface area (SASA; X axis, in $Å^2$), of the two IL-38 ACs and their frequency (Y axis), along a 1 µs aMD simulation of the 3 molecules shown in Figures 3a, 3b and in (D). (F) Recombinant human WT IL-38 2-152 (grey symbols) and a C38S mutant human IL-38 2-152 (black symbols) were treated with $H_2O_2$ (0.1, 1 and 10 mM), or left untreated (lightest grey), and the *in vitro* formation of IL-38 amyloids was assessed by CR staining and fluorescence detection during 18 h at 37 ºC. Both proteins were diluted in a PBS-based buffer for the assay. Results are expressed as mean ± SEM for n=3 independent experiments. A.u., arbitrary units. WT, wild type.

**Fig. S4. Specificity of IL-38 immunostaining of normal human epidermis**

Immunostaining for IL-38 in normal human skin. Representative single plane confocal images are shown from the 3D reconstructed stack in Figure 4a (top row), or from an isotype control stained sample (bottom row). Red staining (second and third columns) represents IL-38 and blue staining (first and third columns) depicts nuclei (DAPI). Dotted lines outline the epidermal-dermal border. Pictures are representative of n=3 different healthy donors. Scale bars, 5 µm.

Figure S1

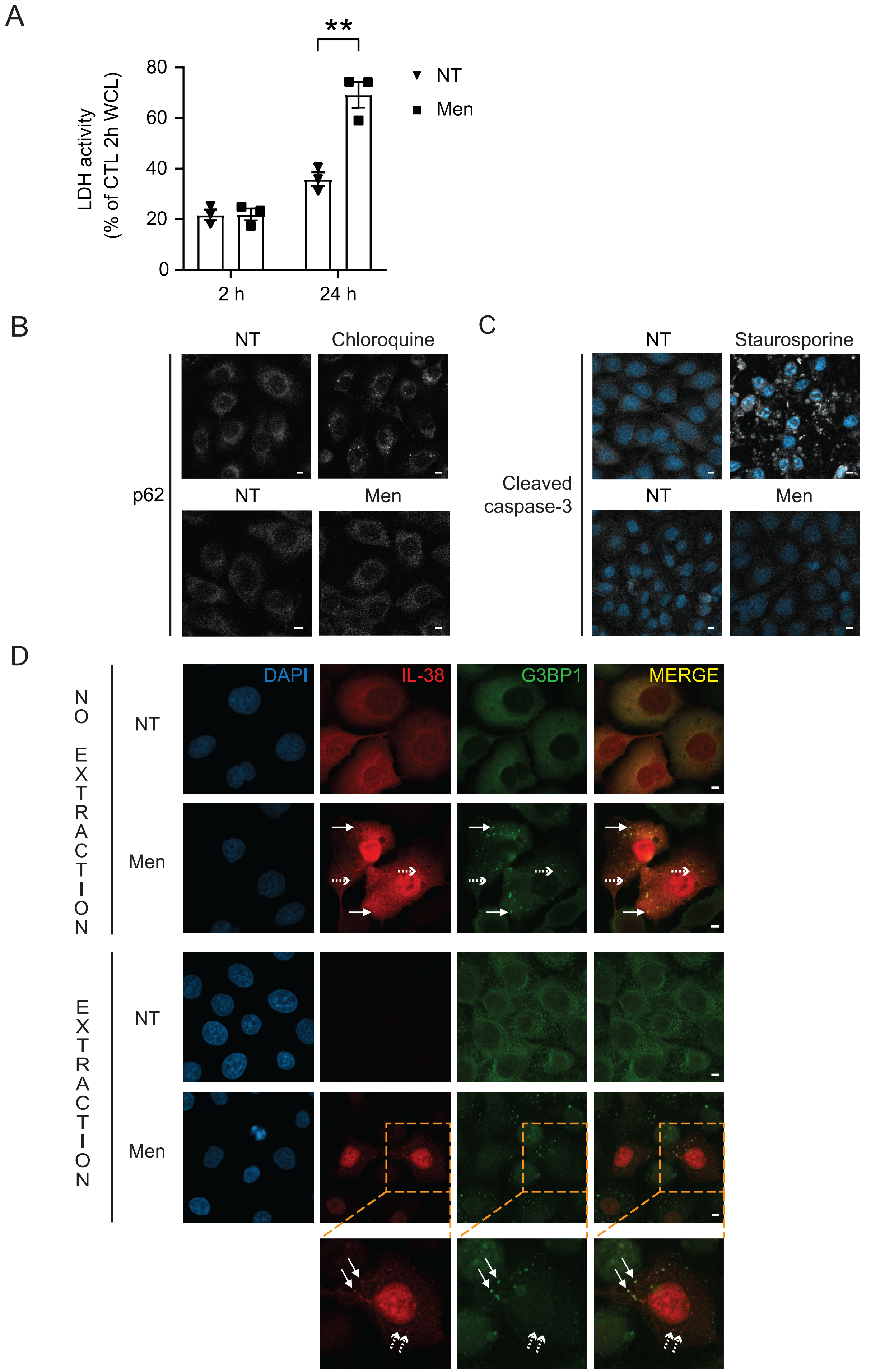
A
**
80
60
40
20
0
LDH activity
(% of CTL 2h WCL)
2 h
24 h
NT
Men
B
NT
Chloroquine
p62
NT
Men
C
NT
Staurosporine
Cleaved
caspase-3
NT
Men
D
NO EXTRACTION
NT
Men
EXTRACTION
NT
Men
DAPI
IL-38
G3BP1
MERGE

### Figure S2

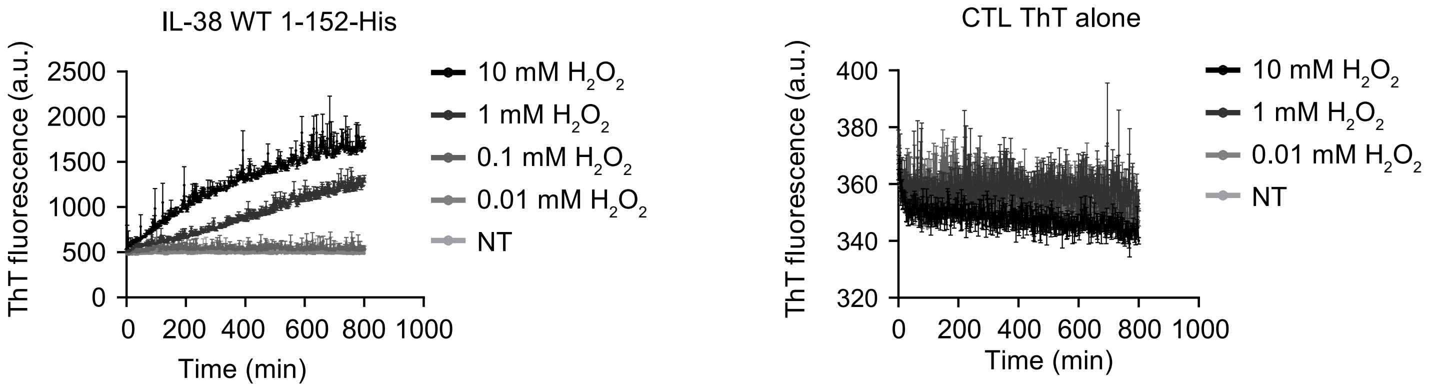
IL-38 WT 1-152-His
ThT fluorescence (a.u.)
2500
2000
1500
1000
500
0
0 200 400 600 800 1000
Time (min)
10 mM H2O2
1 mM H2O2
0.1 mM H2O2
0.01 mM H2O2
NT
CTL ThT alone
ThT fluorescence (a.u.)
400
380
360
340
320
0 200 400 600 800 1000
Time (min)
10 mM H2O2
1 mM H2O2
0.01 mM H2O2
NT

### Figure S3

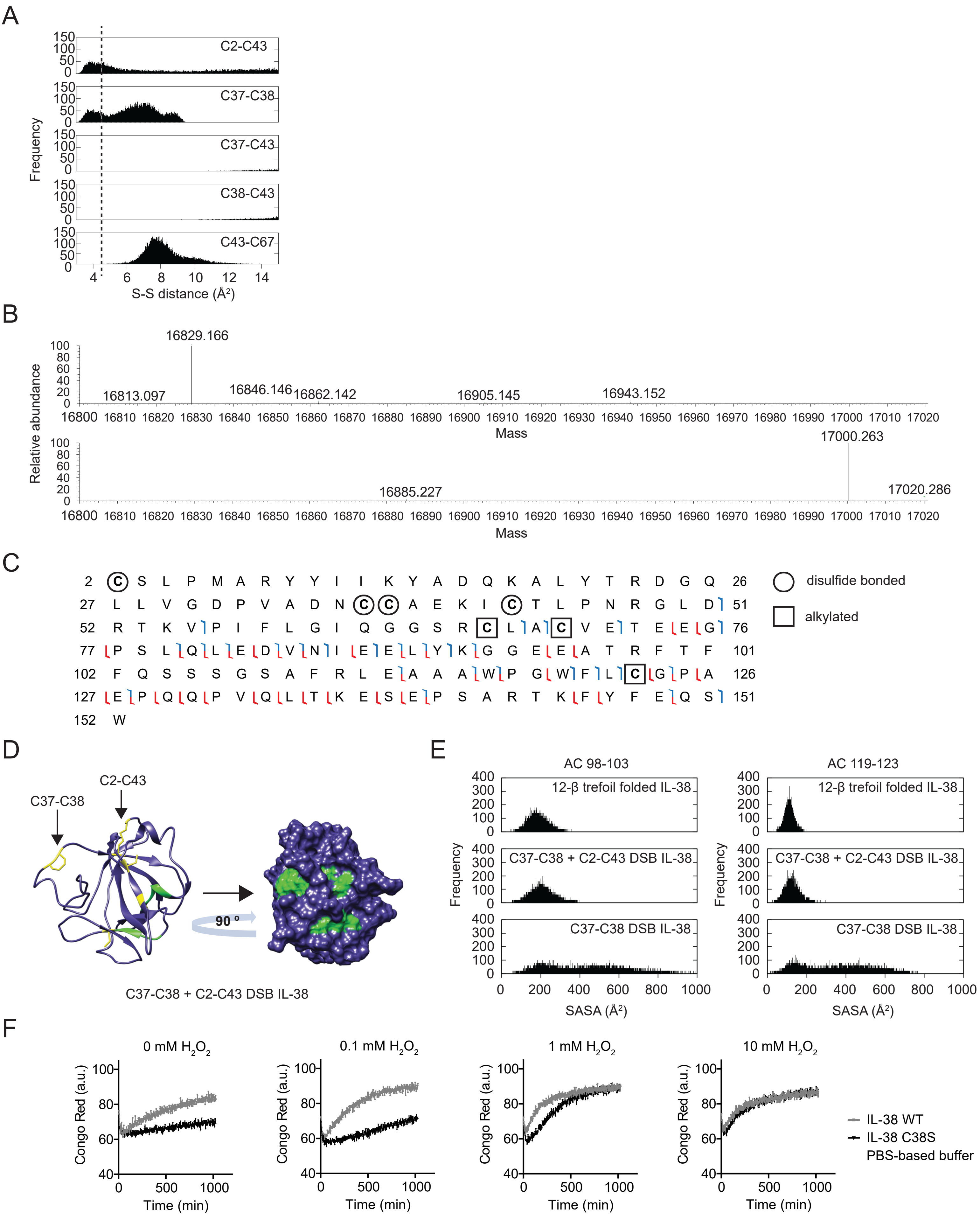
A
Frequency
C2-C43
C37-C38
C37-C43
C38-C43
C43-C67
S-S distance (Å²)
B
Relative abundance
16829.166
16813.097
16846.146
16862.142
16905.145
16943.152
Mass
17000.263
16885.227
17020.286
Mass
C
2 C S L P M A R Y Y I I K Y A D Q K A L Y T R D G Q 26
27 L L V G D P V A D N C C A E K I C T L P N R G L D 51
52 R T K V P I F L G I Q G G S R C L A C V E T E E G 76
77 P S L Q L E D V N I E E L Y K G G E E A T R F T F 101
102 F Q S S S G S A F R L E A A A W P G W F L C G P A 126
127 E P Q Q P V Q L T K E S E P S A R T K F Y F E Q S 151
152 W
disulfide bonded
alkylated
D
C37-C38
C2-C43
90 º
C37-C38 + C2-C43 DSB IL-38
E
AC 98-103
AC 119-123
12-β trefoil folded IL-38
C37-C38 + C2-C43 DSB IL-38
C37-C38 DSB IL-38
Frequency
SASA (Å²)
F
0 mM H₂O₂
0.1 mM H₂O₂
1 mM H₂O₂
10 mM H₂O₂
Congo Red (a.u.)
Time (min)
IL-38 WT
IL-38 C38S
PBS-based buffer

### Figure S4

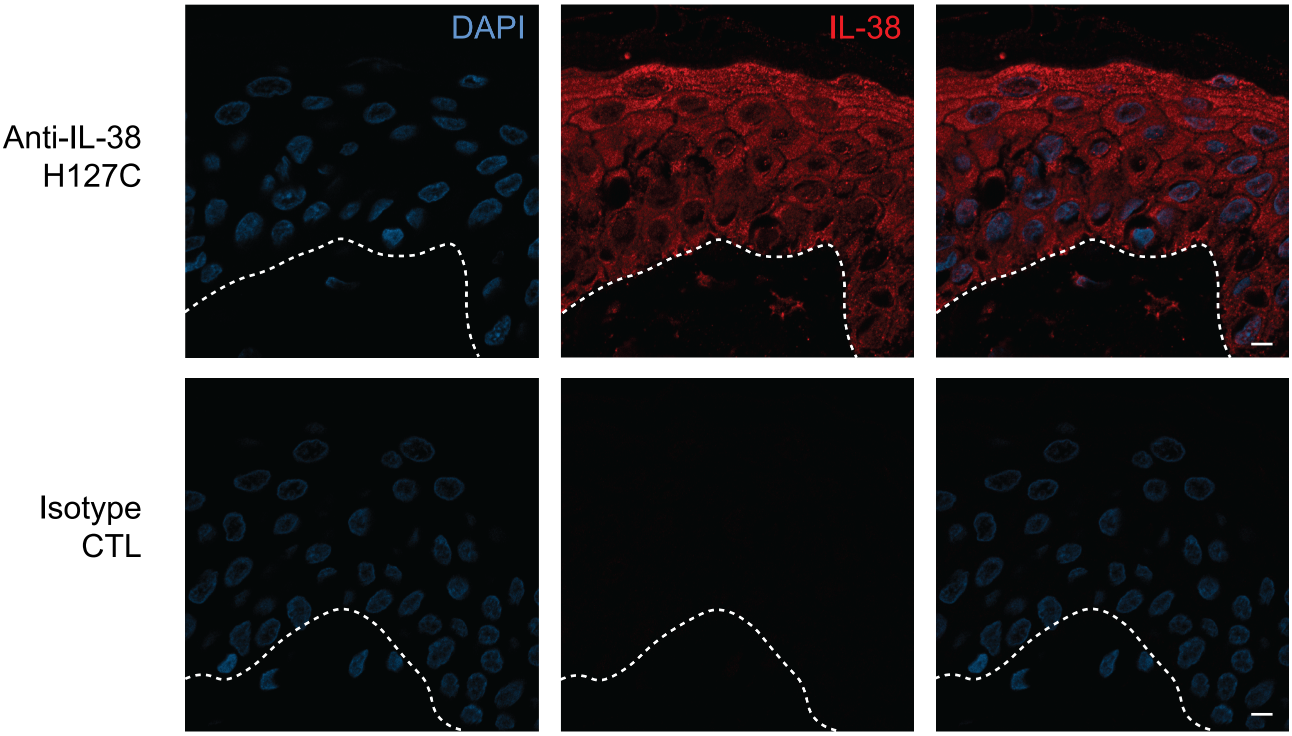
